# Leaderless consensus decision-making determines cooperative transport direction in weaver ants

**DOI:** 10.1101/2023.10.29.564117

**Authors:** Daniele Carlesso, Madelyne Stewardson, Simon Garnier, Ofer Feinerman, Chris R. Reid

## Abstract

Animal groups need to achieve and maintain consensus to minimise conflict among individuals and prevent group fragmentation. An excellent example of a consensus challenge is cooperative transport, where multiple individuals cooperate to move a large item together. This behavior, regularly displayed by ants and humans only, requires individuals to agree on which direction to move in. Unlike humans, ants cannot use verbal communication but most likely rely on private information and/or mechanical forces sensed through the carried item to coordinate their behaviour. Here we investigated how groups of weaver ants achieve consensus during cooperative transport using a tethered-object protocol, where ants had to transport a prey item that was tethered in place with a thin string. This protocol allows the decoupling of the movement of informed ants from that of uninformed individuals. We showed that weaver ants pool together the opinions of all group members to increase their navigational accuracy. We confirmed this result using a symmetry-breaking task, in which we challenged ants with navigating an open-ended corridor. Weaver ants are the first reported ant species to use a ‘wisdom of the crowd’ strategy for cooperative transport, demonstrating that consensus mechanisms may differ according to the ecology of each species.

## INTRODUCTION

Consensus is central to the lives of social animals. Whether they be politicians voting laws or honey bees selecting a nesting site, groups must be able to agree on one option out of mutually exclusive ones (1). Effective decision-making strategies solve conflicts between individuals and, in turn, prevent group fragmentation. Studying information processing and integration within groups is thus central to understanding the dynamics that allow groups to maintain cohesion.

Strategies for achieving consensus may range from following a small minority of influential individuals (‘leaders’), to pooling together the opinions of all group members. Evidence for leadership behaviour has been reported for several animal groups (2), including humans (3, 4), canid packs (5, 6) and bee swarms (7, 8). Leaders may have greater influence on the group’s decisions because they move faster (8, 9) or are larger (10, 11) than their companions, or because they exhibit strong directional preferences (12). The same exists in human groups when individuals are unable to verbally communicate with each other (3, 13). A different but equally widespread strategy is the ‘wisdom of the crowd’ (14–16). It states that, as long as each group member makes an independent assessment of the available options, the accuracy of the group will increase with its size. Indeed, the inaccuracy of each individual’s assessment is largely canceled out by the summing of many opinions. This principle has received extensive support in both human and animal behaviour literature (14, 15, 17–21). These two strategies are best interpreted as the two extremes along a continuum of consensus decision-making mechanisms. Intermediate strategies that combine the two exist, and may be highly effective depending on the context in which animals make decisions.

An excellent example of consensus decision-making is cooperative transport, where two or more individuals carry an item together to a destination. Mostly found in human and ants (22–25), cooperative transport require consensus on which direction to move in. Unlike humans, ants cannot verbally communicate their intentions to the rest of the group. Instead, they rely on chemical signals and mechanical forces sensed through the carried items (23, 25, 26). Furthermore, decisions made by individual ants are based only on locally available information. The decentralised nature of cooperative transport makes it extremely resilient to individual failure, adaptable to environmental conditions, and scalable with group size.

Currently reported in more than 40 ant genera (22, 27), cooperative transport allows colonies to greatly expand the range of prey items that workers can retrieve. In some species, such as *Lasius neoniger* and *Pheidole oxyops*, cooperatively transported items can constitute up to 85% of the total food mass arriving at the colony (28, 29). Cooperative transport is found in ant species that occupy vastly different ecological contexts with distinct environmental challenges. This makes cooperative transport an excellent model system for studying how consensus decision-making may evolve under different environmental pressures. Most commonly, ants encircle the prey item from all directions so that some ants face the desired direction of motion while others face the opposite direction (24). In this situation some ants will have to walk forward, others backwards, and others sideways. Carriers must coordinate their efforts to achieve efficient transport and avoid getting stuck in useless tug-of-war. Several studies have investigated the dynamics underlying cooperative transport in ants (24, 30–35), but how the decisions of each worker are integrated within the group has been seldom explored so far. A notable exception is that of the longhorn crazy ant *Paratrechina longicornis*, which relies on leadership behaviour for navigating during cooperative transport (25, 26).

Comparative approaches for studying decision-making in animal groups are rare. Gelblum, Pinkoviezky (34) proposed a tethered-object approach to investigate the behavioural mechanisms underlying cooperative transport. Here a prey item is tethered to a fixed point using a thin string, and the magnitude of the fluctuations that groups produce around their average direction of movement is measured. This decouples the motion produced by leaders and followers, revealing the consensus strategy used by the group. Groups using a ‘follow-the-leader’ strategy will generate periodic oscillations which increase in magnitude with group size. Large sizes lead to increased behavioral inertia (36) which manifests as deterministic oscillations with higher amplitudes. Groups using a ‘wisdom-of-the-crowd’ strategy will instead produce random fluctuations around the average direction of motion, the magnitude of which is inversely proportional to group size (25, 34). This is because the accuracy of the group will increase with the number of ants engaged with the load. The tethered-object protocol allows us to position ant species along a one-dimensional spectrum of consensus mechanisms, where oscillatory motion is closer to leadership behaviour and fluctuating random patterns are closer to independent pooling of opinions. This simple setup thus represents an excellent comparative approach for studying consensus decision-making across ant species.

Here we used the tethered-object approach to study cooperative transport in the weaver ant *Oecophylla smaragdina*. Weaver ants are a polydomous and arboreal species that inhabits tropical Australasian regions (37, 38), where they are dominant (39, 40). *O. smaragdina* displays an impressive range of cooperative behaviours (40–42), and can integrate visual, magnetic and chemical cues for navigation (43). *Oecophylla* ants are opportunistic predators that feed on arthropods and small vertebrates (40, 44), and they have been observed transporting very large prey items moved by hundreds of cooperating foragers (44). Given the wide range of prey sizes that they can collect, together with their polydomous nesting habit and excellent navigational skills, weaver ants are an attractive system to study cooperative transport.

The aim of our study is twofold. First, we want to investigate the navigational decision-making strategy used by weaver ants to efficiently transport prey items back to their nests. Secondly, we want to establish the tethered-object protocol as comparative tool to study the mechanisms of cooperative transport across ant species. We first challenged ant groups with transporting tethered items to characterise the consensus decision-making strategy they use. We then tested groups in an untethered, symmetry-breaking context, where they had to negotiate their exit from an open-ended corridor to successfully retrieve a prey item back to the nest. In groups using ‘wisdom of the crowd’ strategies, there will be greater conflict of opinion when deciding which side to exit the corridor from, which may induce deadlocking. Outside of the corridor there should be a greater alignment in the directional preference of the group, causing cooperative tranport to be faster, more direct and less likely to stall. Groups that use ‘follow-the-leader’ strategies are predicted to maintain their speed when navigating obstacles but to change direction more often, since they can easily converge to directional consensus (25, 32).

## MATERIALS AND METHODS

### (a) Experimental set-up

Experiments were performed using 9 queenright colonies of the weaver ant *Oecophylla smaragdina* (Fabricius 1775) between December 2021 and December 2022. Colonies were maintained in temperature-controlled rooms at 27 ± 1°C under a 12:12 hours photoperiod at Macquarie University (Sydney, Australia), and fed twice a week with 50% (v/v) sugar water and crickets.

The foraging arena consisted of a white plastic box (64×40×22 cm) with walls coated with talcum powder to prevent escaping of ants (45). The arena was positioned in a metal enclosure (150×83×150 cm) covered with white curtains and evenly illuminated from multiple directions to eliminate orientational cues. The floor of the arena was covered with a paper sheet that was replaced at the end of each testing day (tethered-object experiment) or at the end of each trial (corridor experiment) to remove chemical traces left by ants. The floor of the arena was cleaned with 100% ethanol to remove residual chemical traces at the end of each day. Experimental trials were video recorded using a Panasonic GH4 camera in 4K resolution (3840×2160 pixel) at 24p or 25p.

We used canned grasshoppers (Exo Terra^©^ Grasshoppers XL) as prey items in all experiments. We presented ants with grasshoppers weighing 0.3, 1.3 or 3.5 (± 10%) grams to cause variation in the number of ants carrying the load. Hereafter, we refer to these stimuli as *light*, *medium* and *heavy* items respectively. *Heavy* items were obtained by adding standard weights (magnets, 1g each) inside the grasshopper. *Light* items were obtained by cutting grasshoppers into smaller pieces. *Medium* items were generally obtained using full grasshoppers. All items were weighed at the beginning of each day. We used fresh grasshoppers every day and maintained them at room temperature (20 ± 1 °C) between trials to avoid weight changes due to desiccation.

### (b) Experimental methodology

We starved colonies for 24h before testing to ensure high foraging motivation. On testing days, colonies were connected to the experimental arena through a plastic tube (ø 22 mm). To incentivise recruitment of ants to the arena, we placed a feeder containing 50% sucrose solution and a pinned grasshopper on top of a clean A4 paper sheet in the center of the arena. Ants were allowed to explore the new environment for at least 30 minutes before testing. Trials started when ants successfully formed foraging trails to the food sources. Colonies that failed to form foraging trails within 1 hour were deemed unmotivated and replaced. We removed the A4 paper sheet along with the food sources from the arena before starting experiments.

#### Tethered-object experiment

We tethered a grasshopper to the floor of the arena using a thin 10cm-long string. The grasshopper was initially positioned near its tethering point on top of a clean A4 paper sheet, which ensured no chemical contamination between trials. Ants had to transport the item for 10cm before fully extending the string. We video recorded trials for 10 minutes after ants fully extended the string. At the end of each trial, we removed the A4 paper sheet and the tethered item from the arena. All ants connected to the item were then returned to the colony. Colonies were tested a maximum of 6 times a day with 30 minutes rest between trials. All colonies were presented with each item weight (light, medium, heavy) at least once in pseudo-random order across trials. We performed 10 trials for each item weight (total N = 30).

#### Corridor experiment

We placed a LEGO^©^ corridor (30×5 cm) 25 cm from the nest entry, perpendicular to the ants’ direction of entrance, and positioned a grasshopper at the center of the corridor in a vertical orientation to prevent directional biases (Fig. S8). We then allowed ants to enter the arena and started video recording. Trials lasted until the ants successfully transported the item to the nest entrance, at which point a new trial was set up. Colonies were not allowed to consume the cricket. Colonies that failed to move the item within 45 minutes of the start of the experiment were deemed unmotivated and replaced. We tested colonies a maximum of 5 times a day with 30 minutes rest between trials. The order of presentation of items was pseudo-randomized across trials, ensuring that every colony was presented with each item weight (light, medium, heavy) at least once. We performed 10 trials for each experimental condition (total N = 30).

### (c) Data extraction and analysis

The position of the item was extracted for each frame using a custom computer vision script coded in R (46) (version 4.3.1) using the *Rvision* (47) and *trackR* packages (48). For the tethered-object experiment, the script returned the (x, y) coordinates of the item’s centroid and the angle of its main axis relative to the vertical axis of the image. For the corridor experiment, the script also returned the traffic rate and directionality of ants at both ends of the corridor. All data was scaled using a ruler in frame. The position and number of carriers attached to the item were manually noted every 720 frames. Ants were considered as carriers if their mandibles overlapped with the item without moving for the 10 frames preceding and following each frame of interest. This allowed us to distinguish between ants carrying the item and ants that were transiently sensing it with their mouthparts. Data manipulation was performed in R using the packages *tidyverse* (49), *data.table* (50), *circular* (51), and *trajr* (52).

In the tethered-object experiment, we quantified the fluctuations produced by the group around its average direction of motion for 10 minutes. We calculated the average direction of motion within each trial and assigned it a vector of 0 degrees. This allowed us to compare fluctuations across trials. The angle of deviation from the average direction of motion was then calculated for each frame of the video recordings.

In the corridor experiment, we analysed the trajectory of the item for each trial using the *trajr* package in R. Trajectories within the corridor were analysed from the moment at which the item had moved 2 cm until the moment at which the item exited the obstacle. This allowed us to analyse the conflict between the ants within the corridor without considering initial coordination issues in moving the item. Trajectories outside of the corridor started after ants traveled 3 cm from the end of the corridor – to avoid including data in which ants still differed in their directional opinions – and finished when the group arrived within 3 cm of the nest entrance. These exclusion criteria ensure that any difference in the trajectories’ statistics would be due to the directional opinions of ants rather than other concurrent factors – such as recruitment strategies or insufficient number of carriers. We smoothed all trajectories before analysis to reduce tracking noise, and then calculated the speed, moving speed, number of stops, proportion of path backtracked, and straightness for each trajectory, each defined below.

We considered groups to have stopped when their speed was extremely slow (below 0.1662 mm/s). This threshold was arbitrarily calculated as the 0.15 quantile of the speed distribution across all trajectories within the corridor. The speed and moving speed of the load were calculated as the average speed between each pair of trajectory points, measured in mm/s. Moving speed refers to the speed of the group when removing all segments of the trajectory in which ants were stopped (speed < 0.1662 mm/s). The number of stops was measured as the number of times that the speed of the group fell below 0.1662 mm/s. We standardised the likelihood of groups to stop by dividing the number of stops by the duration of the trajectory, obtaining the number of stops per second of travel. The proportion of path backtracked was calculated as the ratio between the total distance that ants travelled in the direction opposite to the final direction of motion (i.e., if ants escaped the corridor from the right exit, the total distance backtracked is the distance that ants travelled towards the left exit) and the total length of the trajectory. Path straightness was calculated by dividing the total distance travelled by the group by the shortest distance between the initial and final points of the trajectory. Thus, path straightness can only assume numbers between 0 and 1 where 1 indicates a perfectly straight trajectory. Traffic rate and directionality were measured in ants/s at both ends of the corridor.

The distribution of ants around items was obtained by manually noting the coordinates of individuals engaged in cooperative transport. For each frame in which the ants’ coordinates were available, we calculated the directional vector of the group’s movement by extracting the position of the item’s centroid in the next second. We then used this vector to rotate all coordinates so as to align this vector on the horizontal axis, which allowed us to compare the ants’ distribution around items across conditions.

We measured the coordination among carriers using the efficiency index “*R*” (23). *R* represents the rate of delivery as a flow rate per worker, measured in grams-metre per second per worker. We calculated this index as:

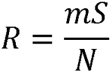

Where *m* is the item weight, *s* is the average speed of groups and *N* is the average group size of ants carrying the item. We used average speed rather than directional velocity so to compare the coordination efforts of groups within and outside of the corridor.

### (d) Statistical analyses

Statistical analyses were performed in R using the *lme4* (53), *glmmTMB* (54), *lmerTest* (55), and *ggeffects* (56) packages. We used generalised linear mixed-effects models (GLMMs) to compare measurements across experimental conditions. Unless otherwise specified, all GLMMs were fit using the *glmmTMB* package. Model diagnostics were performed using the *DHARMa* package (57). Since we wanted to compare the performance of groups across conditions, rather than test the absolute effect of item weight, we included item weight as a categorical predictor in all models. We performed post hoc pairwise comparisons between item weights (*light*–*medium*; *light*–*heavy*; *medium*–*heavy*) and their interaction with position in the arena (within or outside of corridor) using the *emmeans* package (58).

#### Tethered-object experiment

We tested the effect of item weight on carrier group size using a GLMM with Gamma distribution and square-root link. The model included the average number of ants as a dependent variable, item weight as a fixed effect and colony ID as a random effect.

The magnitude of the fluctuations produced by groups was calculated as the range (maximum minus minimum) of angle deviations from their average direction of motion. We compared fluctuations across conditions using a GLMM with Gaussian distribution and log link. We log-transformed the response variable to ensure residual normality and reduce heteroscedasticity. The model thus included the log-transformed magnitude of fluctuations as a response variable, item weight as a fixed effect and colony ID as a random effect.

We calculated the average load carried by ants in each replicate by dividing item weight by the number of carriers. We tested the effect of item weight on the load per ant using a GLMM with Gamma distribution and log link, which included load per ant as a response variable, item weight as a fixed effect and colony ID as a random effect. Item weight was also set as a dispersion parameter to reduce variance heterogeneity.

#### Corridor experiment

We used GLMMs to test whether the average speed, stopping rate, distance backtracked and path straightness shown by ant groups were influenced by their position in the arena (within or outside of the corridor), by item weight and/or by the load transported by each ant. Measurements were averaged over the time in which groups were navigating the corridor and over the time in which ants were outside of it. All GLMMs included item weight and position in the arena as categorical predictors, the interaction between these two variables, load per ant as a continuous predictor and trial ID as random effect. Average group speed was tested using a GLMM with Gamma distribution and log link. Average moving speed was tested using a GLMM with Gamma distribution and log link, which also included item weight as a dispersion parameter to reduce overdispersion. Stopping rates were compared using a GLMM with Tweedie distribution and log link. This distribution is appropriate for modelling non-negative values that include zeros (59). Since both distance backtracked and path straightness of groups are proportions in the closed interval [0, 1], these measurements were compared across conditions using an ordered beta regression in *glmmTMB*.

We tested the effects of item weight and position in the arena on group size and load per ant using GLMMs with Gamma distribution and log link. Models included item weight and position in the arena as fixed effects, the interaction between the two, and trial ID as a random effect. We used a GLMM with Gaussian distribution to test whether the efficiency of groups (“*R*”) was influenced by item weight, position in the arena or the interaction between the two. Trial ID was included as a random effect. We performed pairwise comparisons using Watson’s two-sample test for homogeneity to test whether the distribution of ants around the item’s centroid depended on the groups’ position in the arena – both generally and for each item weight.

## RESULTS

### Tethered-object experiment

We verified that manipulating item weight caused a change in the number of ants engaged in cooperative transport (Fig. 1A). As expected, we found that carrier group size increases with item weight (χ^2^ = 156.41, p < 0.0001). Pairwise comparisons between item weights confirmed significant differences between each condition (*light–medium*: *t* = -7.622, p < 0.0001; *light–heavy*: *t* = -10.805, p < 0.0001; *medium–heavy*: *t* = -3.220, p < 0.001). These results indicate that group size varies as a function of item weight.

**Figure 1.**
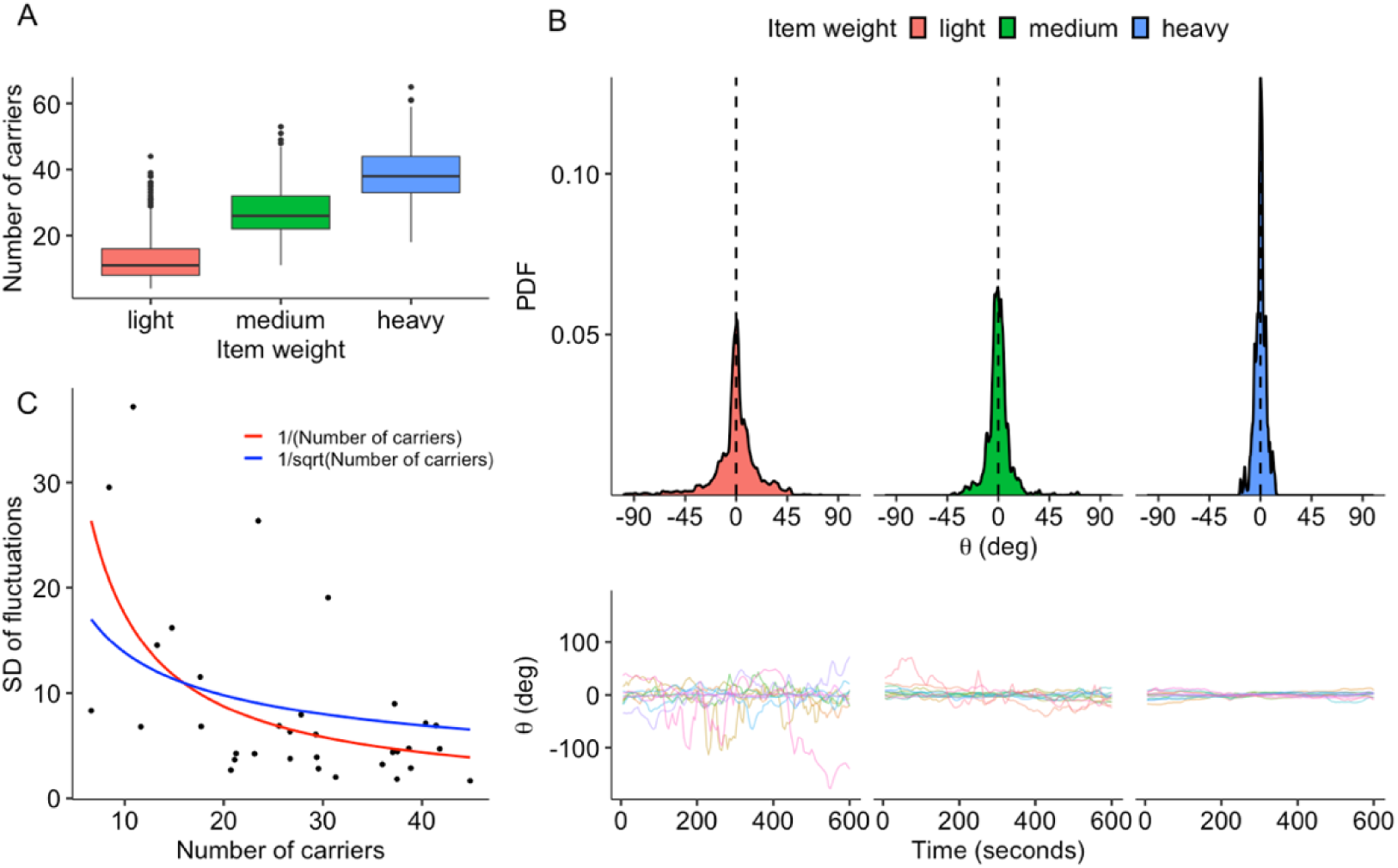
Results of the tethered-object experiment. (A) Boxplot showing median, interquartile range, minimum and maximum values of the number of ants engaged in cooperative transport as a function of item weight. Black dots indicate outliers. (B) Upper panel: Density plot showing the magnitude of fluctuations produced by ant groups as a function of item weight. Black dashed line indicates the average direction of motion of groups. Lower panel: Angle deviations shown by ant groups transporting light, medium and heavy items as function of time. 0 indicates the average direction of motion of groups. (C) Scatter plot showing the relationship between flucutations and number of carriers. Each dot represents one experimental trial. Red and blue lines represent fitted models showing that fluctuations decrease as 1 over the number of carriers (red), and thus faster than 1 over the square root of the number of carriers (blue).

To gain insights on the transport strategy used by ants, we visually analysed the trajectory produced by groups in each trial and found no regular periodicity (Fig. 1B). The magnitude of fluctuations decreased with item weight and, thus, group size (Fig. 1B) (χ^2^ = 33.696, p < 0.0001). Groups transporting *light* items produced larger fluctuations compared to groups transporting *medium* (*t* = 3.326, p = 0.0065) and *heavy* items (*t* = 5.787, p < 0.0001). Accordingly, groups transporting *medium* items showed smaller fluctuations compared to groups transporting *heavy* items (*t* = 2.499, p = 0.047).

We calculated the rate at which fluctuations decrease as a function of group size. For each trial, we extracted the standard deviation of the fluctuations produced by ants and averaged the number of ants transporting the item. We found that the magnitude of fluctuations decreases as one over the number of carriers (Fig. 1C). This result is consistent with the notion of the ‘wisdom of the crowd’, since the noise decays faster than the upperbound of one over the square root of the number of carriers (1/sqrt(N)) that this strategy sets.

We found a positive relationship between item weight and load per ant (χ^2^ = 255.16, p < 0.0001) (Fig. S1). Since the observed decrease in magnitude of fluctuations may also be caused by differences in load per capita, we extracted the fluctuations produced by groups when carrying different item weights but similar load per capita (Fig. S2). This revealed that groups transporting lighter items produced on average larger fluctuations, even when the load per ant is kept constant (Fig. S3). Overall, these results strongly suggest that weaver ants use a ‘wisdom of the crowd’ strategy when cooperatively transporting items.

### Corridor experiment

We extracted the trajectories of ant groups (Fig. S4) and found that ants exited the corridor from the right side in 21 out of 30 trials. The side of exit, however, always corresponded to the side with the higher rate of incoming traffic.

Group speed was higher for lighter weights (χ^2^ = 9.1115, p = 0.0105) and when outside of the corridor (χ^2^ = 70.7911, p < 0.0001) (Fig. 2A). For all item weights, group speed was higher outside of the corridor than within it (*light*: z = -8.414, p <.0001; *medium*: z = -6.276, p < .0001; *heavy*: z = -4.442, p < .0001). Groups were faster when transporting *light* items than when transporting *medium* and *heavy* items both within (*medium*: z = 2.854, p = 0.012; *heavy*: z = 2.480, p = 0.0351) and outside (*medium*: z = 4.324, p < .0001; *heavy*: z = 4.694, p < .0001) the corridor. We found no significant difference in the speed of groups transporting *medium* and *heavy* items within (z = 0.223, p = 0.9729) or outside (z = 1.477, p = 0.3019) the corridor. Group speed increased as the load carried by each ant decreased (z = -3.904, p < 0.0001) (Fig. S5). A nearly-identical pattern in speed was observed when excluding stopping events (Fig. S7).

**Figure 2.**
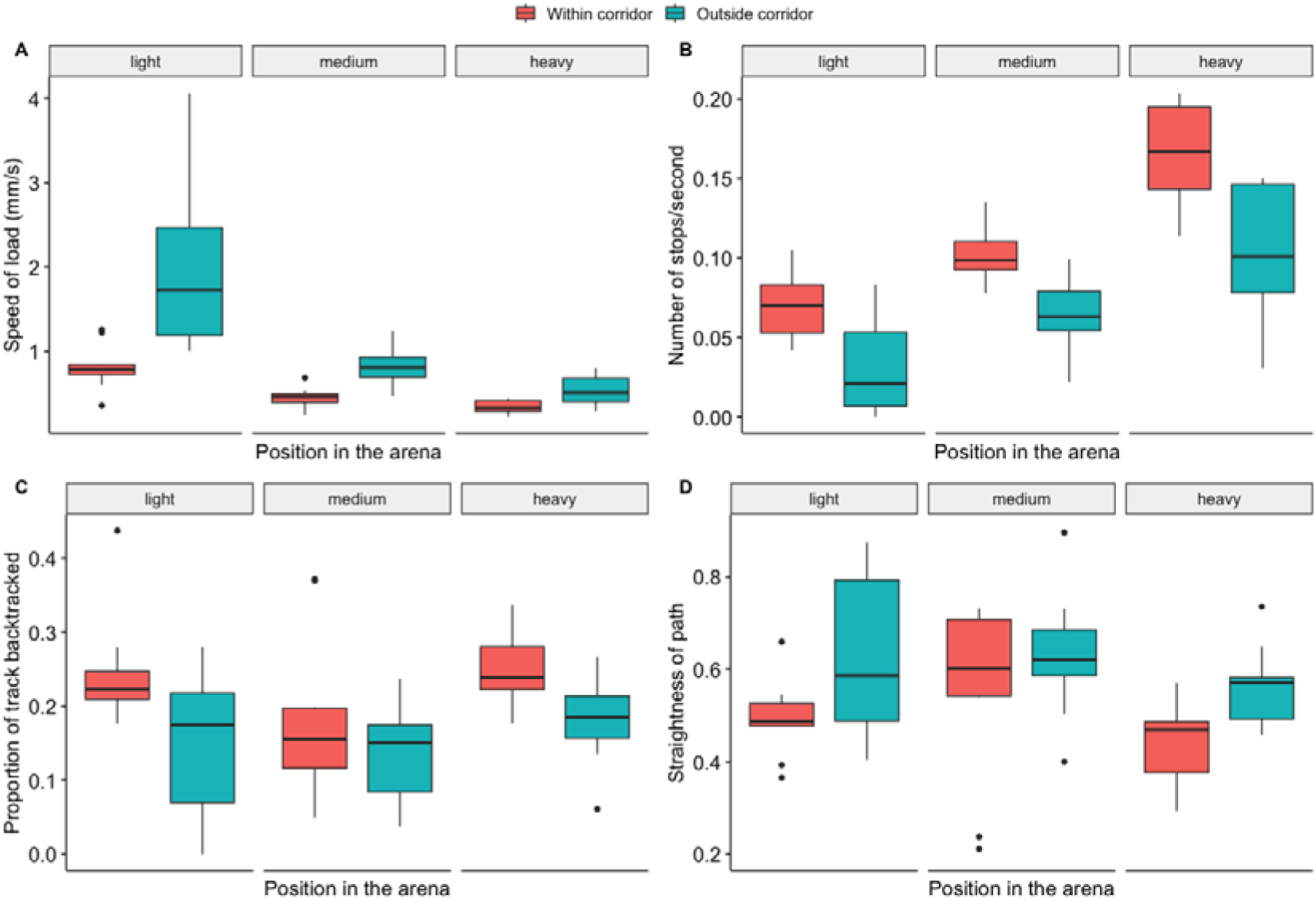
Results of the corridor experiment. Boxplots show median, interquartile range, minimum and maximum values for (A) group speed, (B) stopping rate, (C) proportion of path backtracked, and (D) path straightness as a function of item weight and position in the arena. Black dots represent outliers.

Groups stopped less often when transporting lighter weights (χ^2^ = 13.0533, p = 0.0015) and when outside of the corridor (χ^2^ = 14.5396, p = 0.0002) (Fig. 2B). Stopping rates were lower outside of the corridor for all item weights (*light*: z = 3.813, p = 0.0001; *medium*: z = 2.683, p = 0.0073; *heavy*: z = 3.645, p = 0.0003). Within the corridor, groups transporting *heavy* items showed higher stopping rates than groups carrying *light* (z = -3.465, p = 0.0015) or *medium* items (z = -2.620, p = 0.0239). Although a trend in the same direction can be observed (Fig. 2B), we found no significant difference between the stopping rates of groups transporting *light* and *medium* items within the corridor (z = -1.593, p = 0.2486). Outside of the corridor, ants transporting *light* items stopped less often than ants carrying *medium* (z = -2.922, p = 0.0097) or *heavy* items (z = -4.198, p = 0.0001). A similar trend, although not significant, was found between ant groups transporting *medium* and *heavy* items (z = 1.964, p = 0.1213). Lastly, we found a trend indicating that stopping rates increase with load per capita (χ^2^ = 3.7056, p = 0.0542).

The proportion of path backtracked by ant groups varied with both item weight (χ^2^ = 6.5569, p = 0.0377) and position in the arena (χ^2^ = 7.9104, p = 0.0049) (Fig. 2C). Ants backtracked more within the corridor than outside of it when carrying *light* (z = 2.813, p = 0.0049) or *heavy* items (z = 2.119, p = 0.0341), but not when carrying *medium* items (z = 1.181, p = 0.2374). We found no significant difference between groups carrying different item weights either within (*light*–*medium*: z = 2.220, p = 0.0678; *light*–*heavy*: z = 0.262, p = 0.9628; *medium*– *heavy*: -1.944, p = 0.1264) or outside of the corridor (*light*–*medium*: z = 0.590, p = 0.8254; *light*–*heavy*: z = -0.432, p = 0.9021; *medium*–*heavy*: -1.089, p = 0.5211). Load per ant had no effect (χ^2^ = 0.4418; p = 0.5063).

Ant groups travelled straighter paths outside of the corridor than within it (χ^2^ = 9.1171, p = 0.0025) (Fig. 2D). Path straightness was higher outside of the corridor for groups transporting *light* items (z = -3.019, p = 0.0025), but not for groups transporting *medium* (z = - 1.311, p = 0.19) or *heavy* items (z = -1.790, p = 0.0735). Neither item weight (χ^2^ = 3.8478, p = 0.1460) nor load per ant (χ^2^ = 0.4941, p = 0.4821) were found to have a significant effect. No significant interaction between position in the arena and item weight was detected (χ^2^ = 1.5673, p = 0.4567).

Group size increased with item weight (χ^2^ = 40.9891, p < 0.0001) but not with the position in the arena (χ^2^ = 3.5988, p = 0.0578) (Fig. S6). No significant interaction between the two predictors was found (χ^2^ = 1.5272, p = 0.466). Load per capita increased with item weight (χ^2^ = 45.2895, p < 0.0001) but did not vary depending on the position in the arena (χ^2^ = 3.6970, p = 0.0545) (Fig. 3A). The interaction between these two factors was not significant (χ^2^ = 1.5763, p = 0.4547).

**Figure 3.**
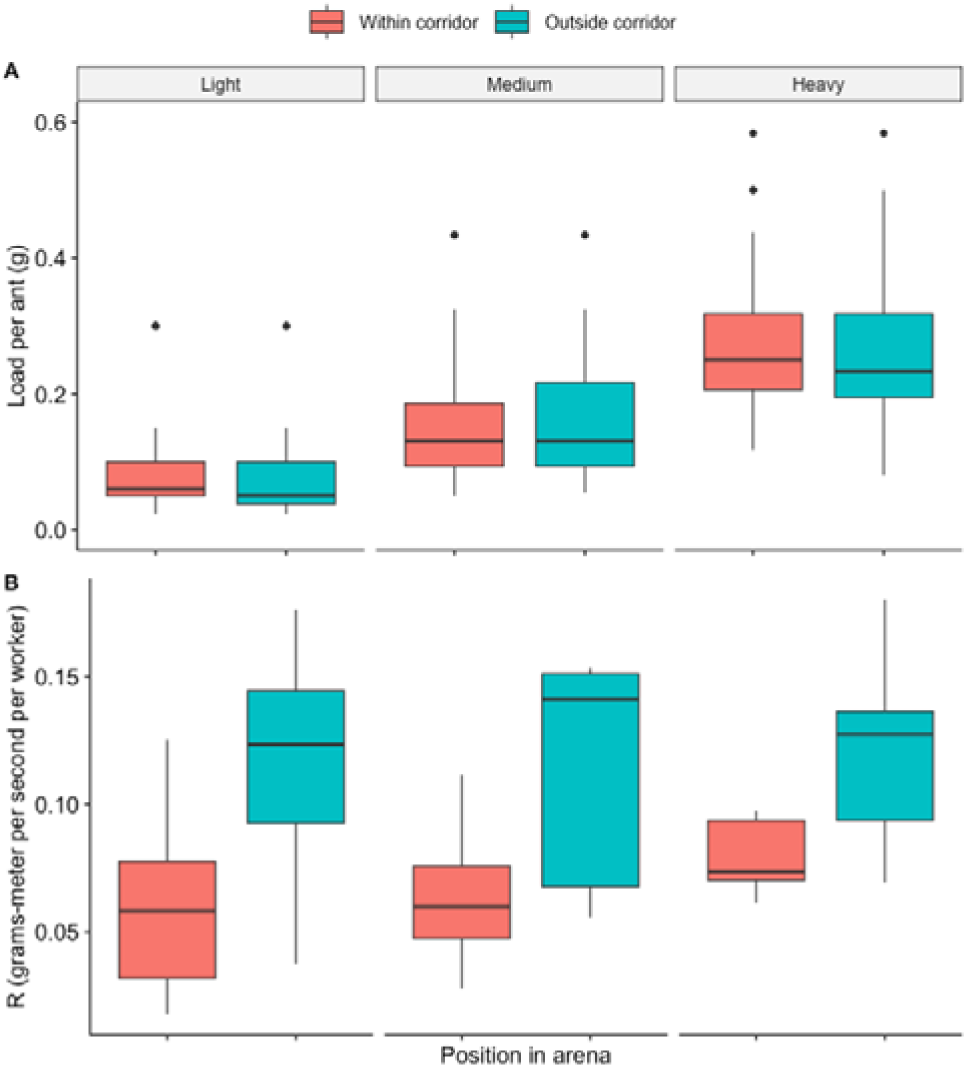
Load per capita (A) and efficiency (B) in the corridor experiment as a function of item weight and position in the arena. Boxplots show the median, interquartile range, maximum and minimum values of the distribution. Black dots show outliers.

Transport efficiency was higher outside the corridor than within it (χ^2^ = 35.1838, p < 0.0001) (Fig. 3B). This relationship was consistent across item weights (*light:* t = -5.932, p < 0.0001; *medium*: t = -5.344, p < 0.0001; *heavy*: t = -4.216; p = 0.0001). Transport efficiency did not vary with item weight (χ^2^ = 1.7833, p = 0.410). Transport efficiency was similar for all item weights when ants were within (*light*–*medium*: t = -0.195, p = 0.979; *light*–*heavy*: t = -1.242, p = 0.435; *medium*–*heavy*: t = -1.047, p = 0.551) and outside (*light*–*medium*: t = 0.174, p = 0.983; *light*–*heavy*: t = -0.165, p = 0.985; *medium*–*heavy*: t = -0.340, p = 0.939) the corridor. The interaction between position in the arena and item weight was not significant (χ^2^ = 1.5204, p = 0.468).

The distribution of ants around the item’s centroid was not affected by the position of the group in the arena when all item weights were pooled together (U = 0.1198, p > 0.10) (Fig. 4). The distribution of ants transporting *light* and *heavy* items differed depending on whether groups were within or outside of the corridor (*light:* U^2^ = 0.457, p < 0.001; *heavy*: U^2^ = 0.287, p < 0.01). No such effect was found for *medium* items (U^2^ = 0.089, p > 0.10) (Fig. S9).

**Figure 4.**
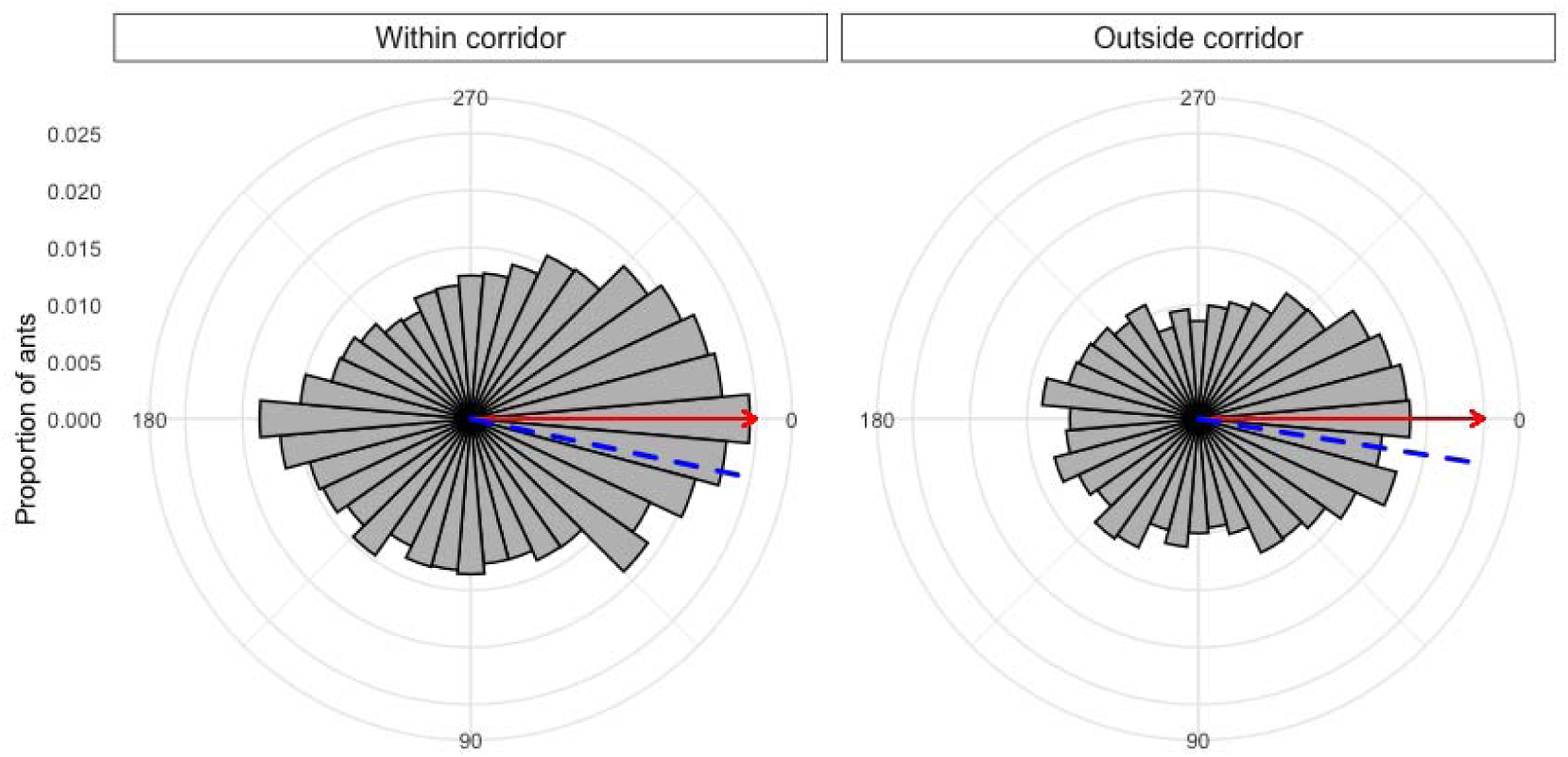
Distribution of ants around the carried item as a function of the position of the group in the arena. Red arrow indicates the direction in which the group is moving, and the blue dashed line indicates the mean of the distribution of ants around the load

## DISCUSSION

Our tethered-object experiment indicated that weaver ants pool together their opinions for navigating during cooperative transport. Groups displayed random nonperiodic fluctuations around their average direction of motion, with magnitude inversely proportional to group size (Fig. 1B – C) consistent with the ‘wisdom of the crowd’ strategy (25, 34). Our findings contrast with the observations made on *P. longicornis* groups, which produced oscillatory patterns that originated from a ‘follow-the-leader’ navigational strategy (25, 34). A potentially confounding element in our experiments is that ants transporting heavier items also experienced higher load per capita (Fig. S1). The decrease in fluctuation magnitude could thus be explained by slower movement, difficulty of transport or other concurrent factors. We ruled this out by showing that the same pattern could be observed when groups transported different item weights while experiencing similar load per capita (Fig. S2). Overall, our results position weaver ants on the opposite end of the consensus continuum from *P. longicornis*.

The ‘wisdom of the crowd’ strategy requires that all ants are roughly informed about the nest position and pull in their preferred direction, without taking into account the forces produced by the other carriers (15). If ants are truly independent from each other, we expect the magnitude of the fluctuations to decrease nonlinearly and faster than one over the square root of group size (1/sqrt(N)). This is because the fluctuations (s.d.) in the force applied by N independent ants into the item should decrease as 1/sqrt(N), but the time scale of these fluctuations might be too fast to be registered by the angular location of the group. In other words, we expect ant groups to switch direction before exhibiting the full 1/sqrt(N) range of locations. The 1/sqrt(N) fluctuations is therefore an upper bound of what the empirical trajectory should show. Here we show that the magnitude of fluctuations produced by groups decreased as 1 over group size (1/N), consistent with an informed-independent model and thus with the notion of the ‘wisdom of the crowd’.

Our symmetry-breaking experiment confirmed our results. Here, ants had to navigate an open-ended corridor in order to retrieve a prey item back to the nest. The presence of a corridor divided the arena in two parts – within and outside of the corridor – varying in the degree of intra-group conflict. We predicted that the higher intra-group conflict would cause groups to be slower, to stop and/or reverse their path more, and to trace more sinuous paths within the corridor than outside of it. All our predictions were confirmed by our results (Fig. 2A – D), and could not be explained by changes in group size or load per capita (Fig. 3A).

The behaviour of *O. smaragdina* and *P. longicornis* groups differ when dealing with obstacles, reflecting the distinct decision-making strategies used by the two species. *P. longicornis* groups confronted with a wall show oscillatory movements that are highly reminiscent of those observed in the tethered-object experiment (25, 34), which ultimately lead ants to escape the obstacle. *O. smaragdina* groups showed no periodic oscillations in our experiments (Fig. 1B). Instead, weaver ants were efficient in maintaining consensus (Fig. 3B) and rarely retraced their steps or stalled for long periods of time. Another important difference between the two species is the dramatic change in speed and stopping rates showed by *O. smaragdina* after exiting the corridor (Fig. 2A – B). Group speed increased on average by 155%, 94% and 56% for light, medium and heavy items respectively. The stopping rate of groups for light, medium and heavy items was 56%, 33% and 41% higher in the corridor than outside of it. These changes cannot be attributed to higher group sizes nor to lower loads per capita (Fig. 3A, Fig. S6), but instead emerge from the reduced intra-group conflict once the corridor is escaped. Conversely, *P. longicornis* groups show little to no changes in speed or stopping rates while navigating obstacles (25, 26, 32, 35). *P. longicornis* and *O. smaragdina* groups thus represent excellent examples of the behavioural consequences of using different consensus strategies in the same context.

We cannot explain the slight rightward bias in our corridor experiment. We designed our experiments to minimise directional biases. Experiments were performed within an enclosure fully covered with white curtains, evenly lit from multiple overhead directions. The corridor was positioned centrally in the arena, and the item placed in the middle of the obstacle in a vertical orientation. We minimised chemical contamination by replacing the substrate and the corridor after each replicate. It is also unlikely that this bias originated from experience or route learning. The bias was evident in the first trials of each experimental day, where the right side was chosen in 8 out of 11 replicates, by ants which had never experienced the arena before. In addition, the time required to navigate the obstacle varied between trials and did not improve in consecutive tests (Fig. S10). In all cases, however, the exit side chosen by ants corresponded to the side of the corridor with the highest traffic rate. The bias may originate from navigational strategies of ants. Side biases are common in animals (60–64). Consistent directional turning and outline tracing has been found in other arboreal ants (65), and it may be a conserved strategy for navigation. A small bias in one direction, combined with the positive feedback of pheromone trails, may aid ants achieving consensus by ensuring asymmetrical traffic around obstacles.

A question that arises naturally from our study is why weaver ants use a ‘wisdom of the crowd’ rather than a ‘follow-the-leader’ strategy. While we cannot provide a definitive answer to this question, this may arise from differences in the information available to *O. smaragdina* and *P. longicornis* workers while engaged in cooperative transport. It has been argued that *P. longicornis* workers carrying items lose access to both vision and smell, invevitably preventing them from navigating (26). Newly attached ants provide groups with updated bearings, transferring environmental information into the system. Weaver ants are exceptional navigators thanks to their ability to integrate visual, magnetic and olfactory cues (43). We occasionally observed carriers tapping on the ground with their antennae (pers. obs.), suggesting that ants may be able to access directional information using pheromone trails, removing the need for temporary leaders to guide the group. Weaver ants also possess large eyes and rely heavily on visual information for navigation (43, 66). They use celestial cues to derive compass information, but can also use artificial light and distal landmarks when necessary. If none of this information is available, ants switch to magnetic cues to orientate. Given their remarkable navigational skills, we hypothesise that weaver ants engaged in cooperative transport possess (at least some) knowledge about the route leading back to their nest.

The consensus strategies used by the two species align with the ecological pressures of their natural habitat. *P. longicornis* ants are opportunistic predators and scavengers that primarily collect proteins from live and dead insects (67, 68). They dominate ecologically thanks to their exceptional navigational skills and short-range recruitment, outcompeting other species by quickly exploiting food resources (67–71). The primary goal of cooperative transport in this species is to retrieve prey items as quickly as possible to avoid intra- and inter-specific competition. The ‘follow-the-leader’ strategy allows them to accomplish this task efficiently, as demonstrated by their ability to navigate obstacles and disordered environments with minimal speed loss (32, 35, 72). Conversely, weaver ants are highly territorial and aggressive predators that prey on live arthropods (37, 73). They are opportunistic scavengers, and can cooperatively transport large preys such as small birds (44). The primary goal of cooperative transport may thus not be rapid movement, as they may instead prefer to attack approaching competitors rather than escaping them (74). The ‘wisdom of the crowd’ mechanism may be more robust for transporting live items, where prey body movements may prevent carriers from sensing the direction in which other ants are pulling. Being able to withstand and actively generate forces more than 100 times their own body weight (75, 76), *O. smaragdina* is much stronger than *P. longicornis* and among the strongest ants in the world. This may favor slower solutions that rely on brute force rather than faster strategies requiring high coordination among individuals. Future studies on cooperative transport in other species should investigate the links between consensus mechanisms and ecological pressures. For instance, *Pheidole pallidula* ants generate an oscillatory motion when tested in a tethered-object task - indicating a ‘follow-the-leader’ strategy (25) – and share similar ecological pressures to *P. longicornis* (77, 78).

Comparative tools are rare in behavioural ecology, because experimental approaches must be adapted to the species under study. One aim of our study was to enstablish the tethered-object approach as a comparative tool for investigating cooperative transport across ant species (25). Cooperative transport has been reported in more than 40 ant genera (22, 27), but quantitative studies are limited to only a handful of species (24, 25, 29, 30, 79). Even when considering the available research, results are difficult to compare because of differences in methodology. The tethered-object protocol offers an easy-to-run and cost-effective solution to this issue. This protocol simulates the presence of an obstacle, allowing the observations of the movement of ant groups over long periods of time (34). This approach can be readily used in laboratory and field conditions without the need for advanced equipment. In fact, this setup only requires a prey item, a thin string, an anchor point and a camera. Results can be obtained relatively quickly by tracking the movement of the group over time and by estimating the number of ants engaged in transport. Further, there is no need for detailed analyses on the individual-level behaviour of ants, which are often difficult or lengthy to obtain. This approach has proven effective for studying cooperative transport in *P. longicornis* and *Pheidole pallidula* (25), and now in *O. smaragdina*. We hope that the current study will motivate researchers in applying this protocol to other ant species. The tethered-object protocol is limited in that it only describes a linear space of consensus models, delimited at either end by the ‘wisdom of the crowd’ and ‘follow the leader’ strategies. Its results are useful to determine which strategy ants are closer to, but not for identifying possible intermediate strategies and their consequences. For instance, carriers could be only partially informed about the nest direction but also partially coupled with the movements of other ants. Further studies are necessary to investigate in detail intermediate navigational consensus strategies and their advantages compared to known alternatives. Given its simplicity and affordability, we believe that the tethered-object protocol is a valid comparative tool to characterise the navigational strategies of ants during cooperative transport. For the same reasons, this protocol is also an excellent candidate for citizen science and educational events (80).

Here we showed that weaver ants pool together their opinions for achieving navigational consensus during cooperative transport. This allows them to efficiently move along their route and escape obstacles without getting stuck in lengthy deadlocks. The ‘wisdom of the crowd’ strategy has been reported in several animal species (14, 81, 82), humans (14, 83) and even bacteria (84). Our findings adds to the previous literature demonstrating the effectiveness of this strategy in maintaining consensus during collective navigation tasks. A wide survey of cooperative transport is now needed to reveal the relative spread of consensus strategies among ant species and the links with their ecology.

## Acknowlegdements

D.C. thanks Zoe Korzy Wild for help in proofreading the manuscript. D.C. thanks Macquarie University and the Australian Research Council for financial support. D.C. was funded by an Australian Research Council Discovery Early Career Research Award (to C.R.R, DE190101513). C.R.R. was funded by an Australian Research Council Future Fellowship (FT220100669). S.G.’s contribution was supported by the National Science Foundation under Grant No. 1955210 and No. 2222418. O.F. was supported by the European Research Council under the European Unions Horizon 2020 research and innovation program (Grant Agreement No. 770964).

