## Supplementary Materials for "Leaderless consensus decision-making determines cooperative transport direction in weaver ants"

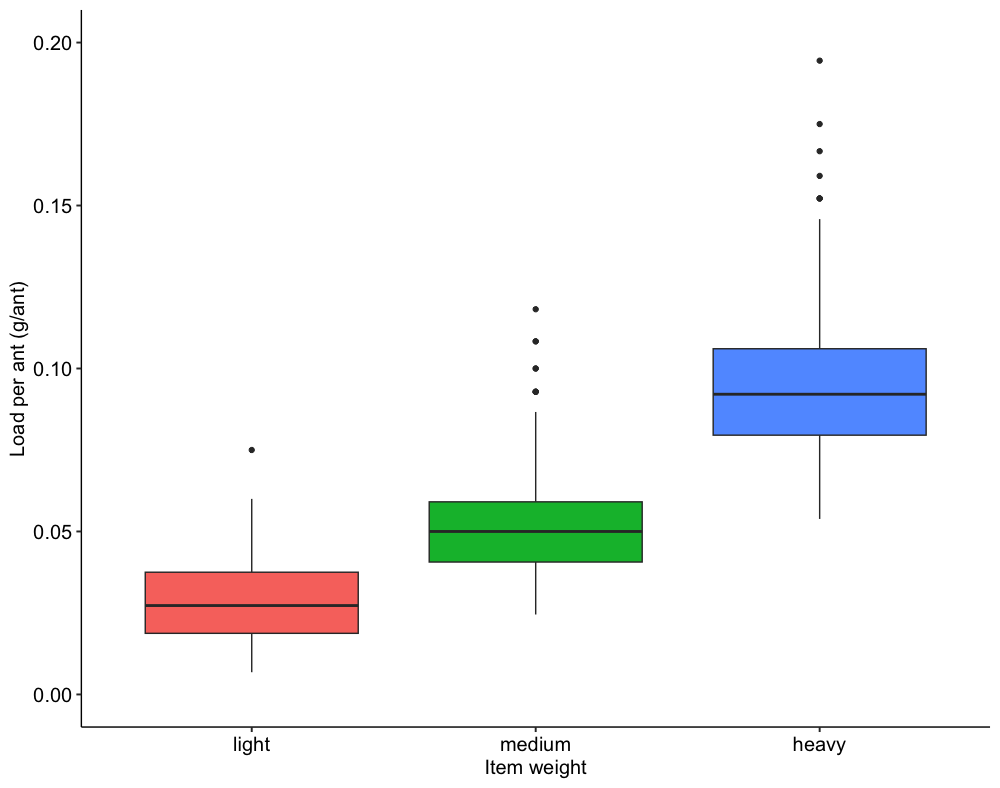

Figure S1 – Load per ant as a function of item weight in the tethered-object experiment. Boxplot shows median, interquartile range, minumum and maximum values of the distribution. Dark dots represent outliers.

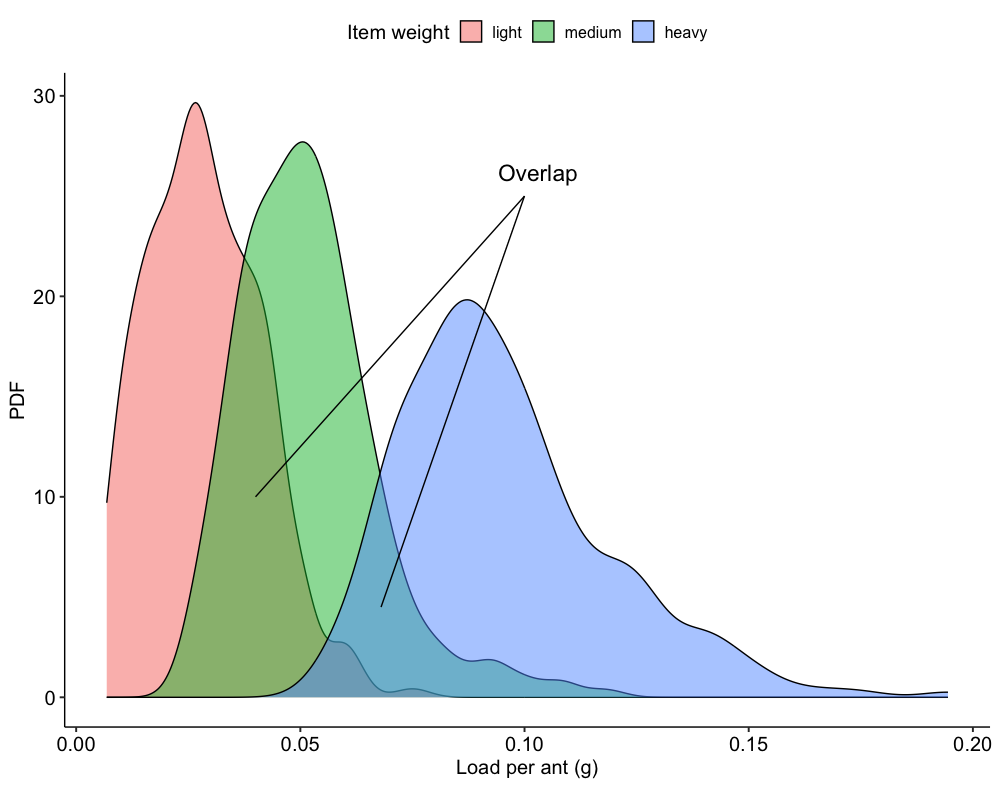

Figure S2 – Density plot showing the distribution of load per capita carried by groups for different load weights in the tethered experiment. Black arrows indicate areas of the distribution where ants transporting different item weights carry similar load per capita.

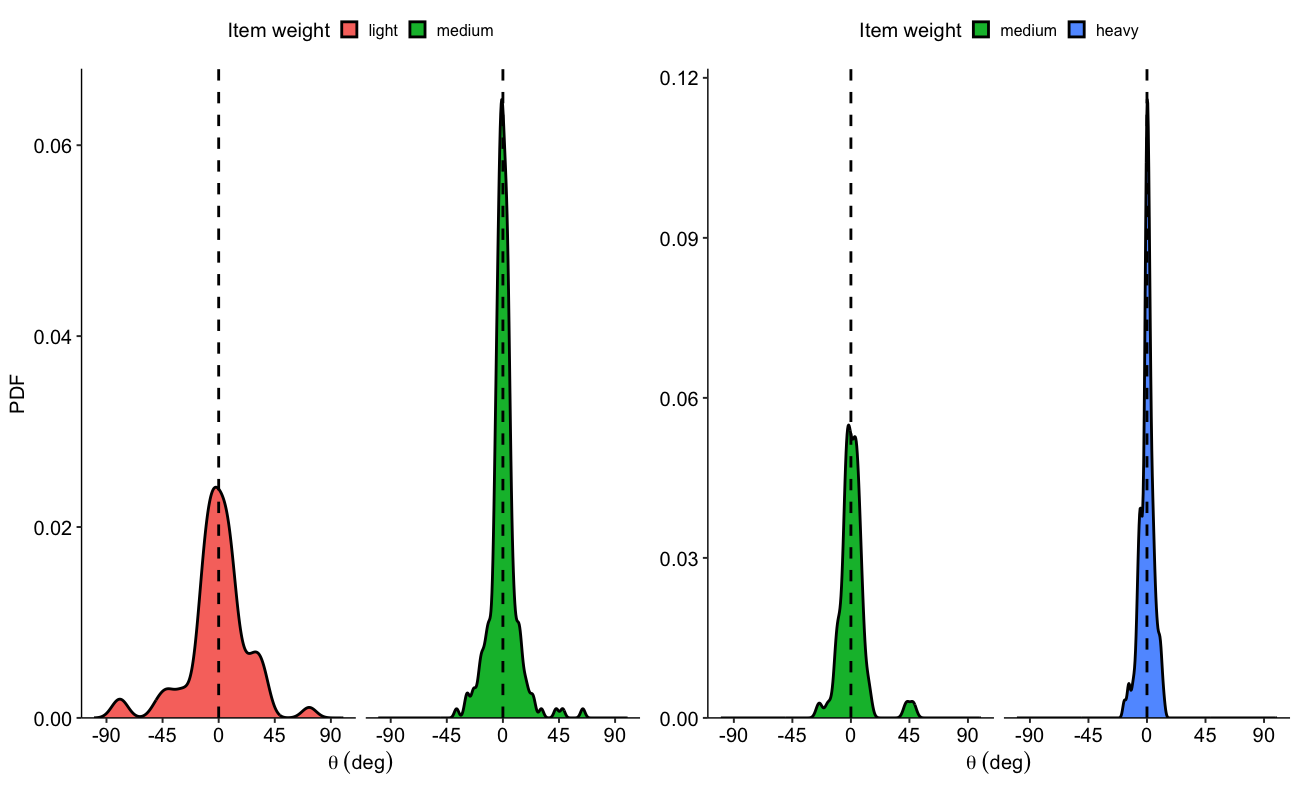

Figure S3 – Density plot showing the magnitude of fluctuations produced by ant groups as a function of item weight. Black dashed line indicates angle (θ ) of 0, which represents the average direction of motion of groups.

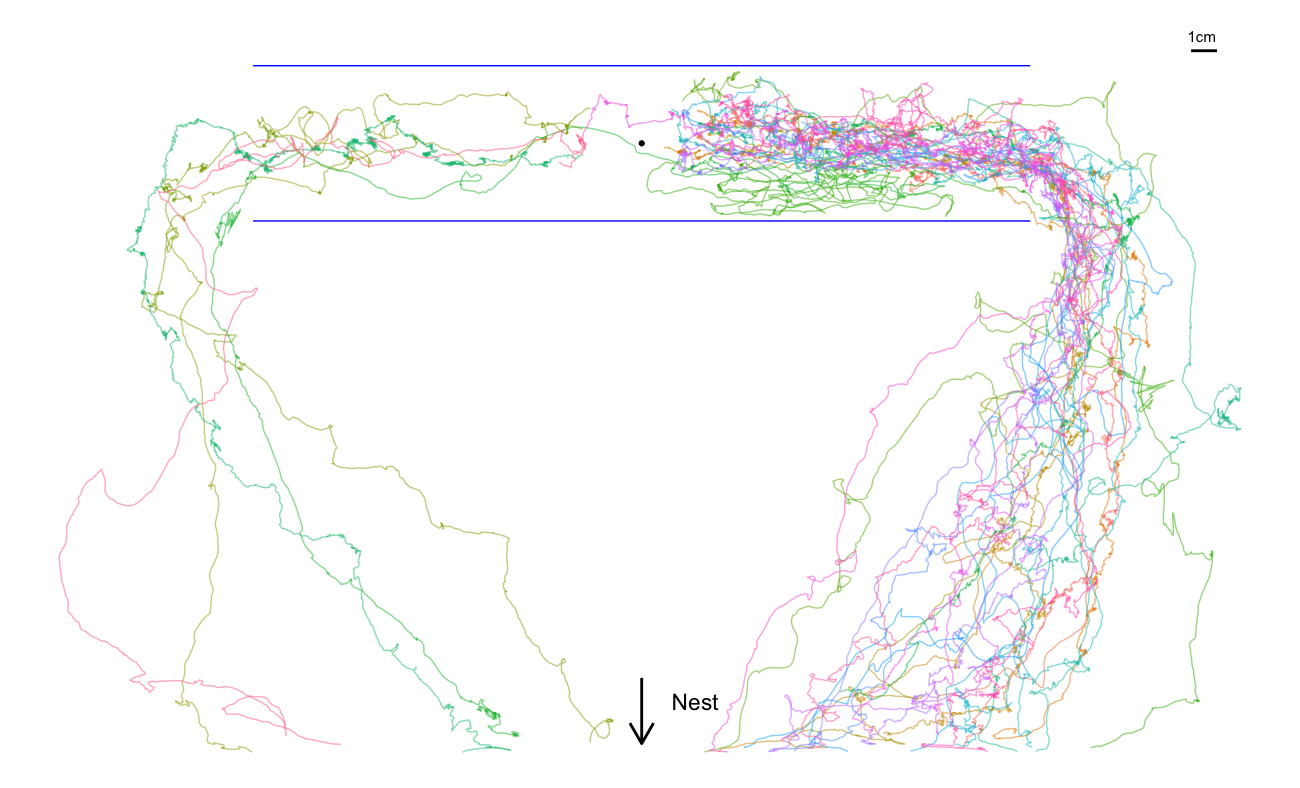

Figure S4 – Trajectories of ant groups in the corridor experiment, after removing the first two centimeters of groups’ movement. Solid blue lines show the walls of the corridor, black dot indicates starting point of the item.

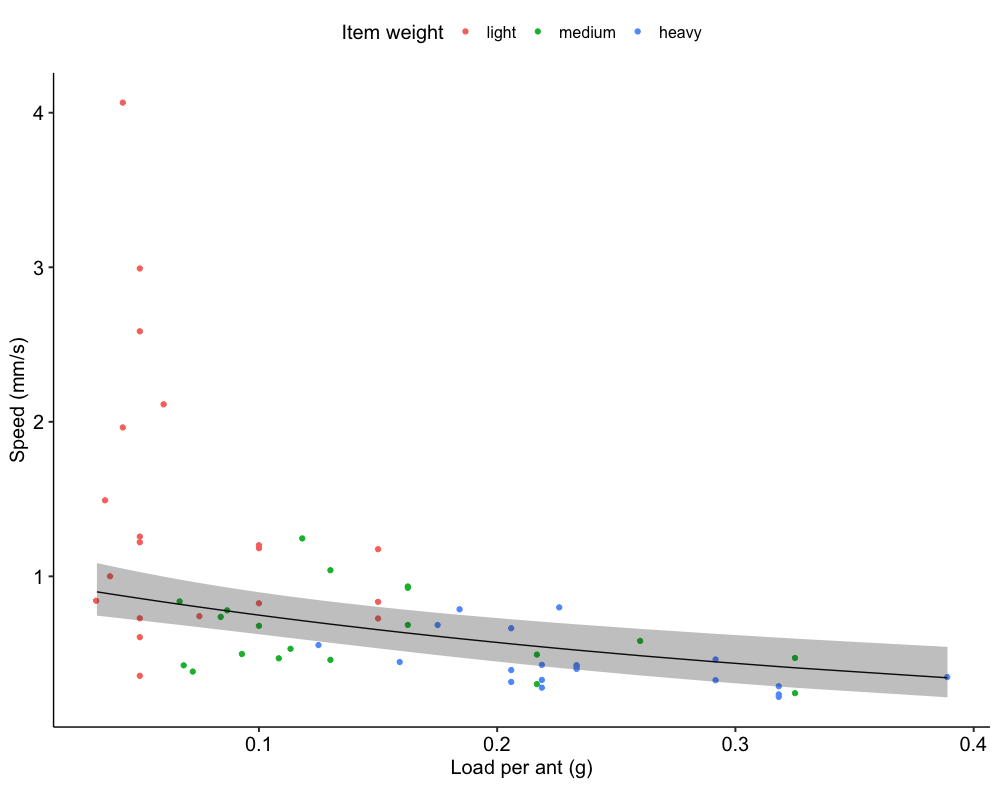

Figure S5 – Speed of ant groups increases as the load per ant decreases. Dots of different colors indicate different item weights. Black line shows prediction of GLMM while grey ribbon shows 95% confidence interval.

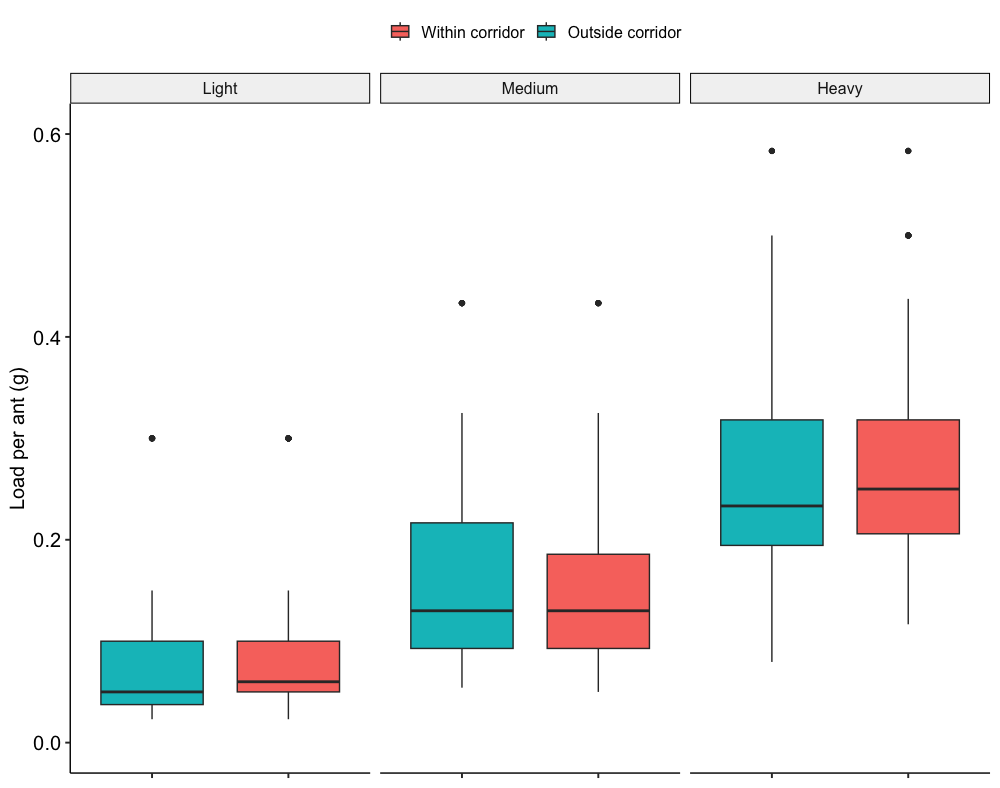

Figure S6 – Number of ants engaged in cooperative transport in the corridor experiment, as a function of item weight and position in the arena. Boxplot shows median, interquantile range, minumum and maximum values of the distribution. Dark dots represent outliers.

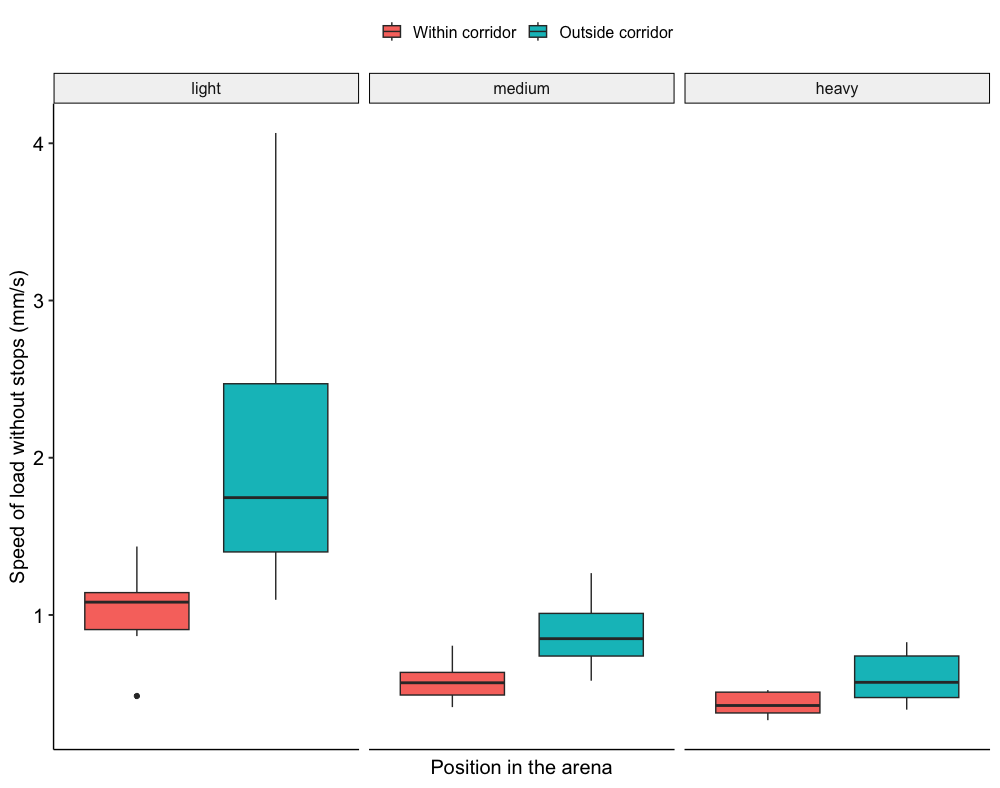

Figure S7 – Speed of groups as a function of item weight and position in the arena. Boxplot shows median, interquantile range, minumum and maximum values of the distribution. Dark dots represent outliers.

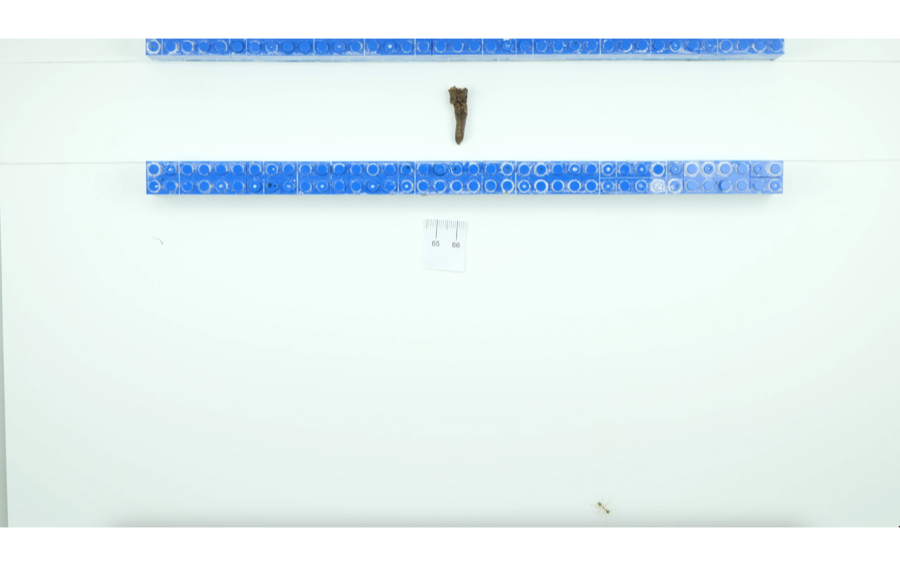

Figure S8 – Experimental arena and initial setup for the corridor experiment.

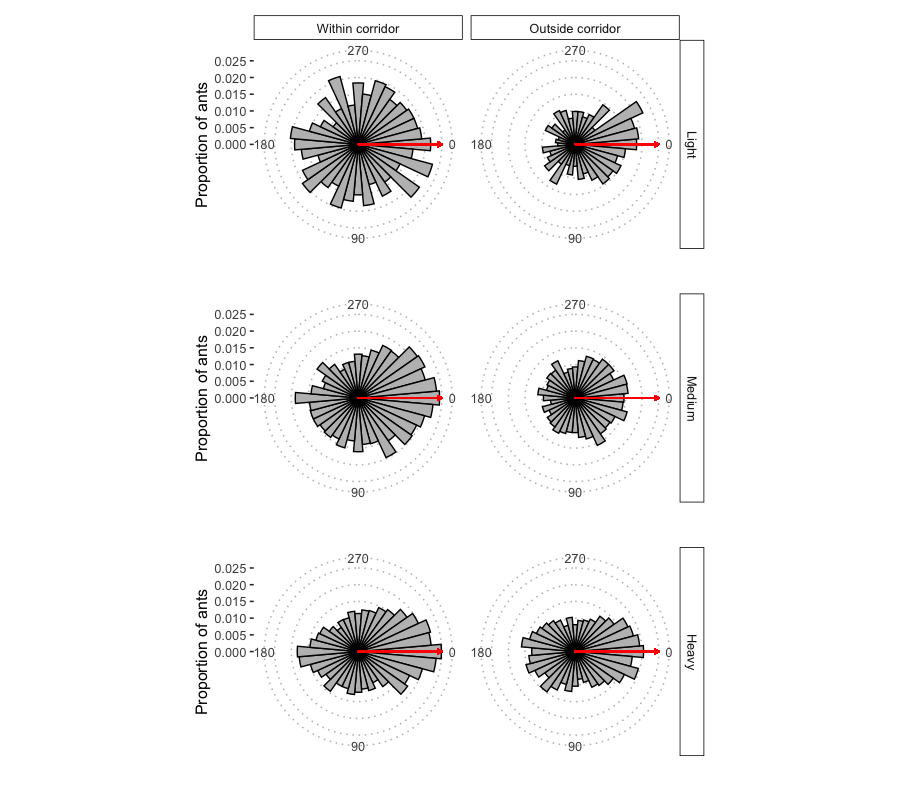

Figure S9 – Distribution of ants around the carried item as a function of position in the arena (columns) and item weight (rows). Red arrow indicates direction of motion of the groups.

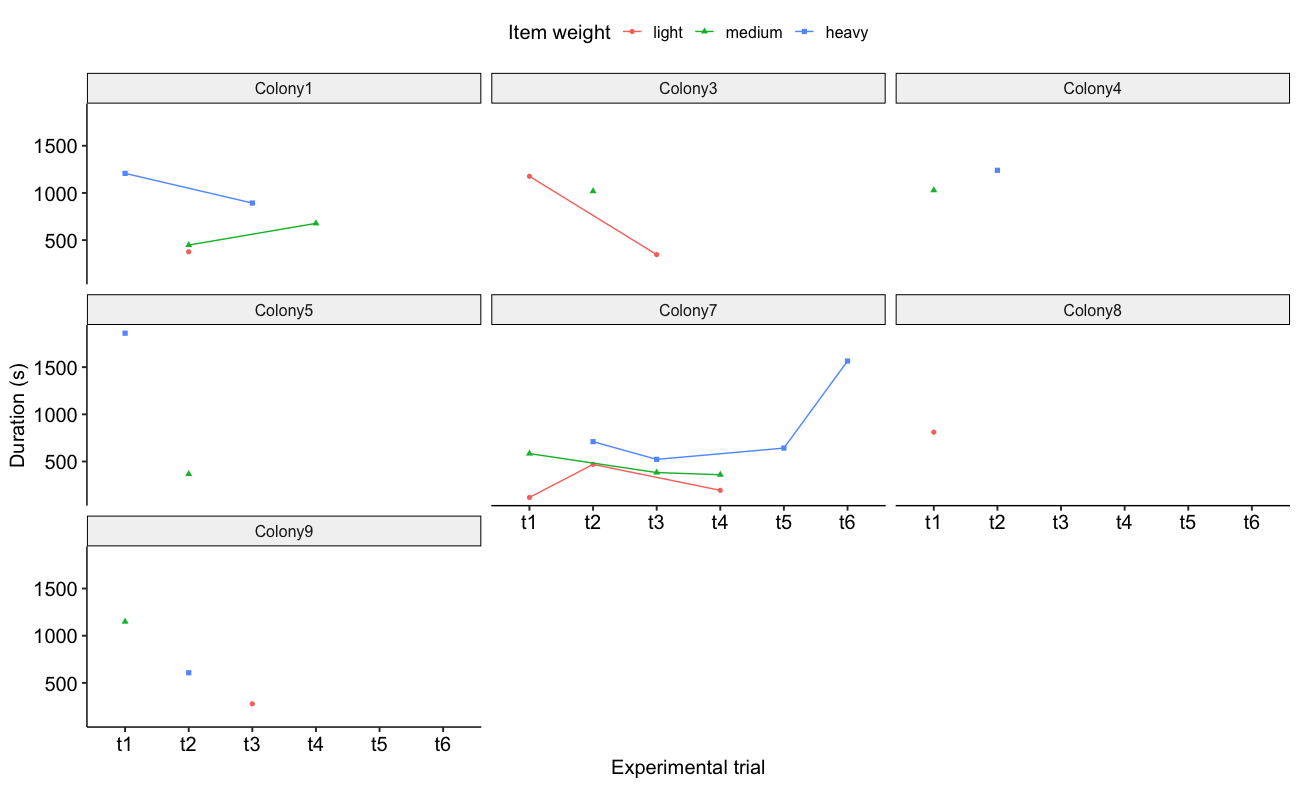

Figure S10 – Time taken by groups of ants to exit the corridor as a function of experimental trial. Color and shape of dots indicate different item weights. Lines connect replicates with the same item weight for ease of visualisation.
